# A survey of biosecurity measures applied on dairy cattle farms in Spain

**DOI:** 10.1101/673996

**Authors:** Francisco Javier Villaamil, Ignacio Arnaiz, Alberto Allepuz, Mikel Molins, Mercedes Lázaro, Bibiana Benavides, Sebastián Jesús Moya, Jordi Casal, Eduardo Yus, Francisco Javier Diéguez

**Author notes:** Corresponding author, (FJD).

## Abstract

Attention to biosecurity has been highlighted as the most important measure to reduce and prevent the introduction of diseases to farms. There is little published information about the biosecurity of dairy cattle in Spain. We therefore aimed to assess and characterize the current application of biosecurity measures on dairy cattle farms in Spain, and relate these to bovine viral diarrhea and infectious bovine rhinotracheitis. From July 2017 to April 2018, data on biosecurity measures for 124 dairy herds were collected using a questionnaire. We also assessed the sanitary status of these farms (efficacy of measures implemented against both diseases using antibody ELISA. Data were analyzed descriptively, and using multiple correspondence analysis and a two-step cluster analysis. Measures to prevent disease introduction were often poorly implemented. Three main clusters of farms were identified: Clusters 1 and 2 included herds of small and intermediate sizes, respectively. These, particularly cluster 1, showed the most deficiencies in the control of vehicles and visitors. However, individual purchases usually involved low numbers of animals, especially in cluster 2, and animals were tested for bovine viral diarrhea and infectious bovine rhinotracheitis at their places of origin or on arrival at farms. Farms in clusters 1 and 2 were frequently under voluntary control programs. Cluster 3 had the largest herd sizes, with somewhat better biosecurity control of vehicles and visitors. However, farms in this cluster also purchased the most animals, sometimes without testing, and hired external workers most often. Farms in cluster 1 showed the best sanitary level, followed by clusters 2 and 3. Collecting data such as these is an important first step to identification of biosecurity shortcomings, and to structuring of adequate follow-up to ensure that measures are implemented correctly on farms in Spain.

## Introduction

Infectious agents that affect livestock may be transmitted by various routes such as live infected animals, trucks and other vehicles, people, aerosols, fomites, or wildlife or insect vectors. Thereby, prevention through biosecurity is the most cost-effective protection (1).

Within the context of animal production, biosecurity is defined as management activities that reduce the opportunities for infectious agents to gain access to, or spread within, a production unit (2).Thus, it has two main components: external and internal biosecurity. External biosecurity entails preventive measures and risk reduction strategies designed to avoid the introduction of pathogenic infections (hazards), whereas internal biosecurity entails measures to limit within-farm transmission of infectious hazards between animals (3).

The importance of biosecurity is highlighted in the European Union health strategy. From 2007 onwards, the EU embraced a new motto as part of its Animal Health Strategy: “prevention is better than cure”, now implemented by the European Commission. The aim was to focus on preventive measures, disease surveillance, controls, and research to reduce the incidence of animal diseases and minimize the impact of outbreaks (4).

The putative benefits of undertaking biosecurity for disease prevention and/or control include improvements in production efficiency (hence greater profits), animal welfare, immune responses to vaccines, and job satisfaction for producers, herd health professionals, and other agricultural workers (5). In addition, in pig herds, a link between biosecurity and antimicrobial treatment-related criteria has been demonstrated and quantified (6).

Despite these benefits, implementation of biosecurity on dairy cattle farms is often sub-optimal, and poor or inappropriate knowledge transfer is often cited as a potential cause of disease spread (7).The main limitations and strengths of the biosecurity measures applied in dairy cattle farms have been studied recently in several countries (7–10), as a preparatory step to develop greater awareness of the importance of each measure and the factors that might restrict its application. In Spain, despite the economic importance of milk production in some regions, current biosecurity practices on dairy farms have not been studied empirically although a recent study assessed perceptions and practices applied by rural veterinarians (11).

The sanitary status of dairy cattle farms with respect to endemic diseases such as bovine viral diarrhea (BVD) and infectious bovine rhinotracheitis (IBR) is highly variable among Spanish regions. In some, the situation is unknown and the approach to outbreaks depends mainly on the actions of individual veterinarians and farmers. In others, mainly in the northwest, voluntary programs were established some time ago, and now involve a large proportion of the cattle population (12).

BVD and IBR, both caused by the BVD virus (BVDV), and bovine herpesvirus 1 (BoHV-1) are highly contagious diseases of economic and trade importance for the livestock industry worldwide. They can cause serious economic losses, with pathogenic effects including reduced milk yield, infertility, abortion, respiratory disease, and an increase in opportunistic infections such as calf pneumonia and scours (13,14). For all these reasons, several European countries have implemented official voluntary or compulsory programs to eradicate both diseases (15,16).

The aim of the present paper was to assess and characterize the current application of biosecurity measures on dairy cattle farms in Spain, and the relationship of these to BVD and IBR.

## Materials and methods

### Area description and herds surveyed

We conducted our study in two Autonomous Communities (AC) of Spain: Galicia (NW), and Catalonia (NE). Galicia is the main dairy cattle area of the country, with 55% of the farms and 38% of the milk production. The mean herd size per farm is 43 cows, lower than the national average of 59.3, and farms are still predominantly family owned and managed. This region is representative of the production type prevailing in the NW and Cantabric area of the country. In contrasts, farms in Catalonia have a mean herd size of 144 cows, with 4% of the farms nationally yielding 11% of the milk production (17). The farm typology in this region could be considered representative of the rest of Spain.

We selected124 dairy farms, 93 from Galicia and 31 from Catalonia, as part of a larger national project on risk analysis for the introduction of BVDV and BoHV- 1todairy cattle herds. For farm selection, the project was presented to the veterinarians responsible for health management in the main dairy cattle areas from both regions. Subsequently, together with the veterinarians, the project was communicated to farmers and those interested in participation were enrolled in the study.

### Biosecurity questionnaire

Data were obtained using a questionnaire which was completed during personal interviews with farmers and the veterinarian responsible for health management of each farm. Farms were visited between July 2017 and April 2018.

The questionnaire, which is available in Spanish on request from the authors, included four sections: I) general farm information (e.g. location, herd size, vaccination programs); II) animal movements (e.g. origin of the animals, frequency of introductions, test, quarantine facilities, external rearing farms, cattle fairs or competitions, pasture) and neighborhood (i.e. other ruminant farms in a 1 km radius); III) movements and types of vehicles and equipment (for live and dead animal transport, manure, slurry and feeding vehicles, machinery or materials) and biosecurity-related measures (e.g. vehicles may enter inside the farm perimeter, vehicles may enter with other animals.) and IV) visitors and staff (e.g. external workers, frequency of professional or nonprofessional visitors that may contact animals, use of protective clothing).

### Health status

The BVDV and BoHV-1 sanitary status of the farms was determined using antibody ELISA, taking into consideration the fact that farms that applied vaccines used inactivated vaccines in case of BVDV and marker vaccines (live or inactivated) in case of BoHV-1.

For BVDV, antibodies against the p80 antigen were determined using a commercial blocking ELISA(BVD p80 Ab, IDEXX laboratories, the Netherlands), since the antibodies of animals vaccinated with inactivated vaccines react mainly with structural proteins rather than the p125 or p80 antigens (18). Additionally, in Galicia, when tested samples indicated possible persistent infection (PI) in an animal (i.e., when a positive result was obtained fora young heifer), this was confirmed by antigen capture ELISA (IDEXX BVDV Ag Serum Plus, IDEXX Laboratories; the Netherlands) based on detection of the E^rns^ viral protein. Two samples positive for BVDV from the same animal taken 3–4 weeks apart were considered to confirm persistent infection.

For the case of BoHV-1, two different tests were used depending on whether vaccines were used on the herd (IDEXX IBR gE antibody, IDEXX Laboratories; the Netherlands) or not (IDEXX IBR gB antibody, IDEXX Laboratories; the Netherlands). All analyses were performed following the recommendations of the manufacturer.

Using the results of these tests, three different BoHV-1 and BVDV farm profiles were established: (1) farms with recent or active infection (BVDV seropositivity in heifers from 9 to 24 months born on the farm and/or PI confirmed), (2) farms with seropositive adult animals but all rearing heifers (9-24 month) seronegative, or(3) free farms (all animals tested seronegative).

### Statistical analysis

All statistical tests were performed with SPSS 15.0. Initially, the frequencies of the different BoHV-1 and BVDV profiles and biosecurity measures were analyzed.

After that, to characterize the current application of biosecurity measures in dairy cattle farms from Spain, a multiple correspondence analysis (MCA) was performed. MCA aims to reduce a set of possibly correlated variables (including all the biosecurity variables and the sanitary status of the farms) to a smaller group of linearly uncorrelated dimensions. We set the number of dimensions to two, to facilitate two-dimensional graphical representation. The position of the full set of categories for each investigated variable (category-points) on the MCA graph is the basis for revealing relationships among variables: variable categories with a similar profile tend to group together, whereas negatively correlated categories are located on opposite sides of the graph. In addition, a two-step cluster analysis (TSCA) was performed to identify clusters of farmers with similar biosecurity levels and BVDV and BoHV-1 profiles. For this analysis, the frequencies of entry of animals, vehicles, and visits (collected as numerical values) were processed as categorical (4 categories based on the quartiles of the variables), but finally were transferred to the results as mean and median frequencies of entry.

## Results

BVDV and BoHV-1 profiles of the124 farms are summarized in Table 1. The serologically free farm category represented similar proportion of the total for both viruses (50% and 44.3% for BoHV-1 and BVDV, respectively). However, the proportion of farms categorized as recent/active infection was much higher for BVDV (36.1% *versus* 10.7%). Thirty-seven of 124 farms used inactivated vaccines against BVDV, whereas the remainder did not vaccinate; 33 used marker vaccines for BoHV-1 whereas the remainder did not vaccinate.

**Table 1.**
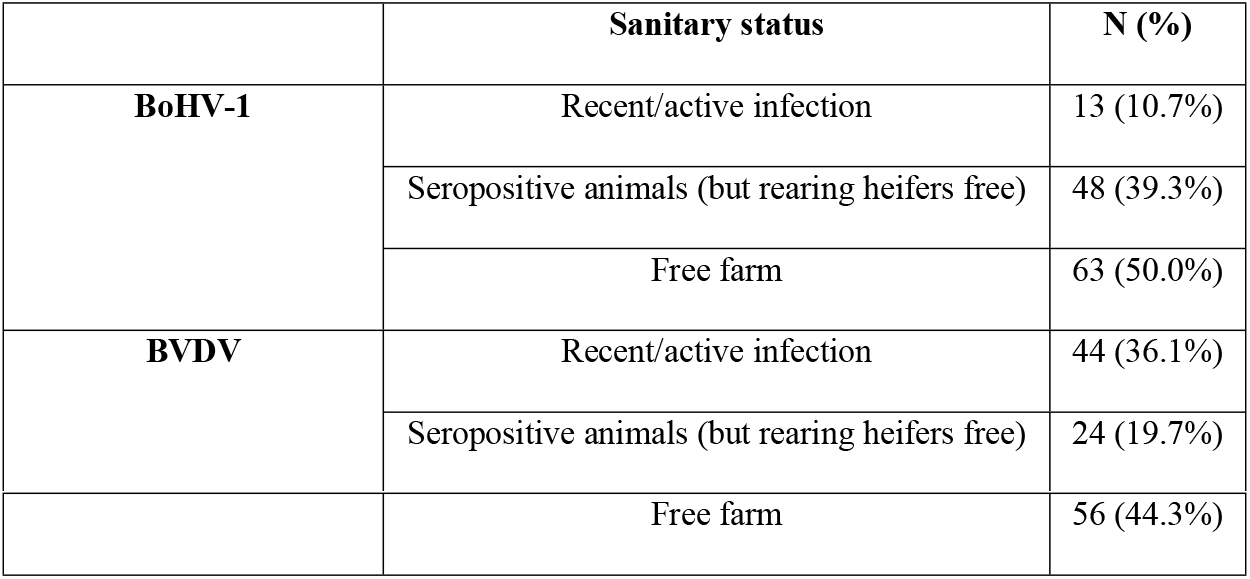
BVDV and BHV-1 sanitary status of 124 farms in Spain

Table 2 describes biosecurity measures related to animal movements and possible contact with other domestic ruminants. For most farms, the risk of introduction of diseases through the purchase of animals (heifers or cows) was presumably negligible, as most reared their own replacements. Farms that purchased cattle (46/124, 37.1%) had a low mean frequency of introductions (5.2 purchased animals/year). However, it is noteworthy that 11/46 (23.9%) farms purchased animals without any testing, 39/46 farms (84.8%) purchased cattle that could contact other cattle during transport, and most farms lacked adequate quarantine facilities. Animal movement to cattle fairs or competitions was a possible pathway of disease transmission for only 7/124 farms (5.7%), but it is noteworthy that nearly all farms reported that animals returned to the farm without any quarantine. Movements to pasture were much more frequent: 54/124 farms (43.6%) reported this type of movement, and half of these reported the possibility of contact with other domestic ruminants at the pasture. Moreover, most farms had other cattle or sheep/goat farms within a 1 km radius.

**Table 2.**
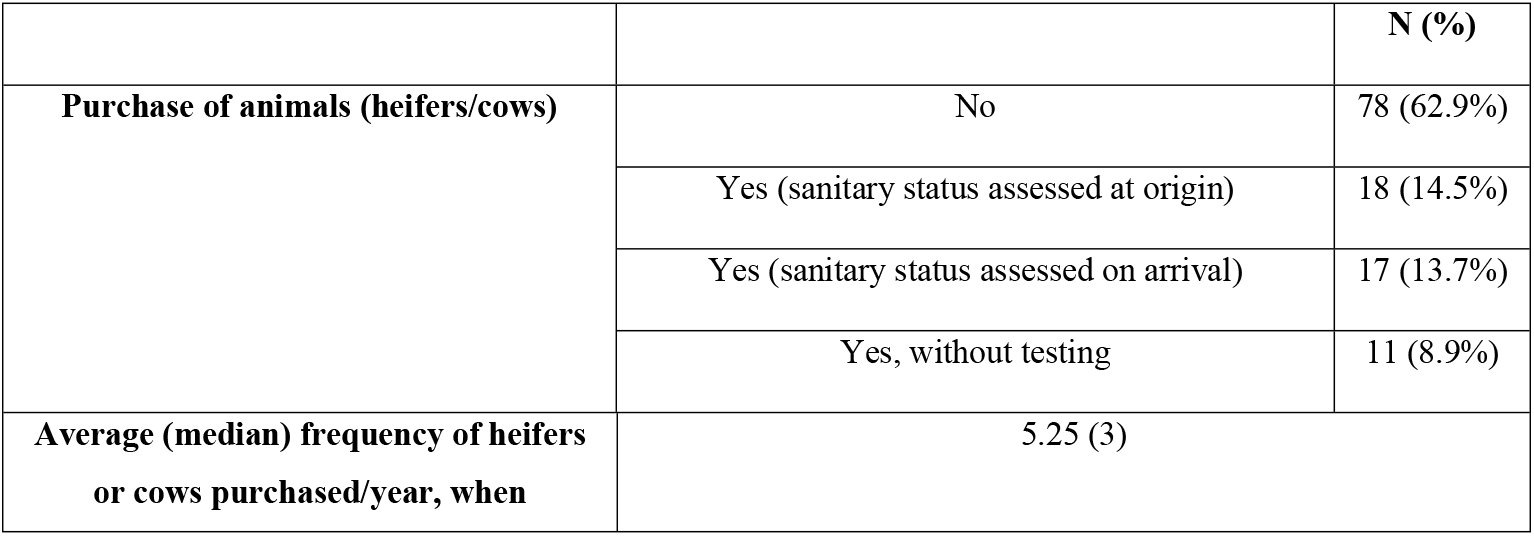
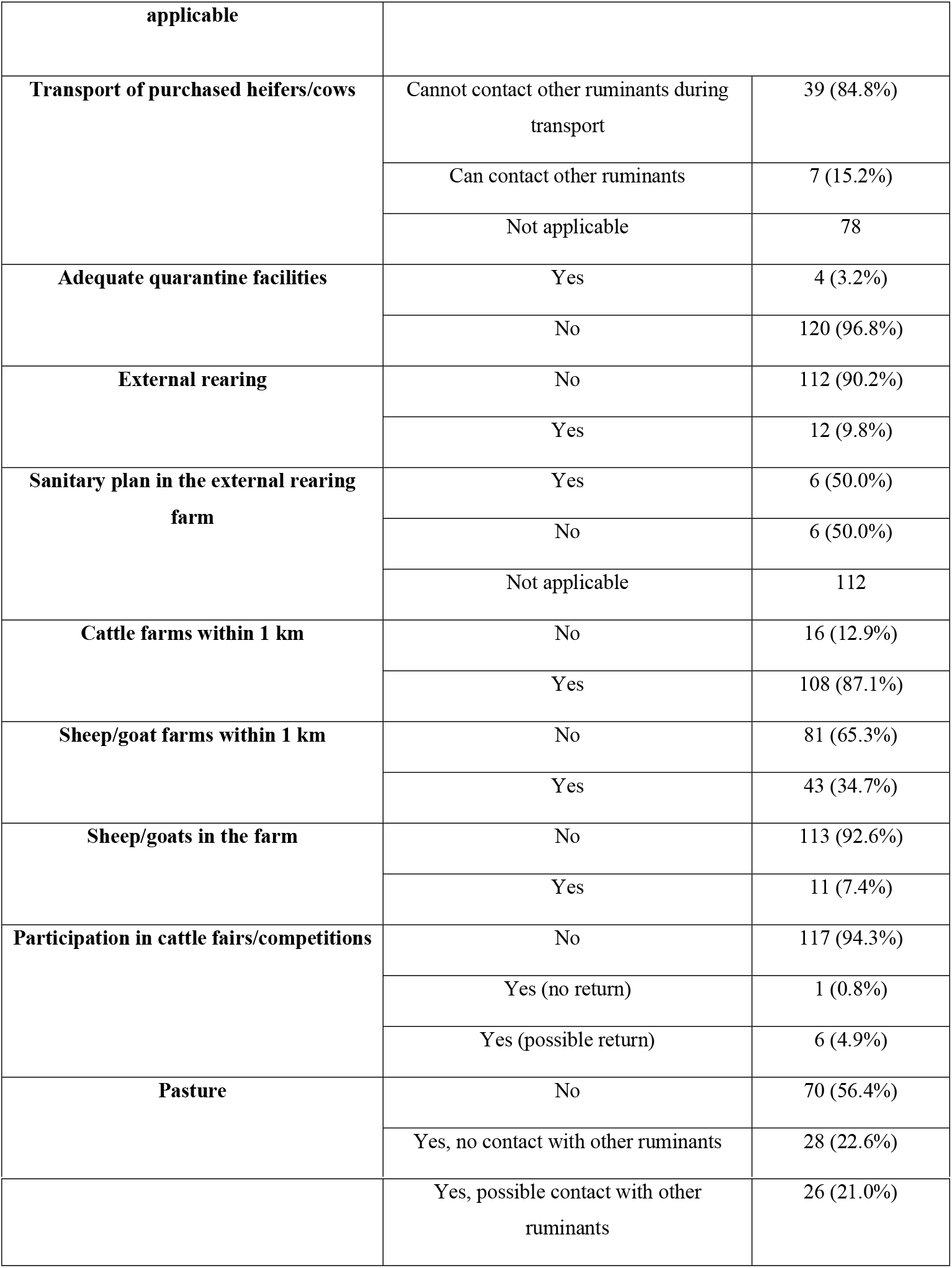
Biosecurity measures related to purchase of cattle, or possible contact with other ruminants, for 124 farms in Spain

Vehicles visiting the farms represented an infection risk for most of the surveyed farms, because nearly all vehicles entered inside the perimeter of all farms (Table 3). The most frequently entering vehicles were feeder wagons, followed by those collecting animals for slaughter, or calves for feedlot. Moreover, in most farms (92.6%), vehicles in the latter two categories were permitted to arrive carrying domestic ruminants from other farms. Several farms shared machinery or other vehicles, such as manure or slurry vehicles, with other cattle farms.

**Table 3.** Biosecurity measures related to vehicles and equipment in 124 farms in Spain

|  |  | N (%) |
| --- | --- | --- |
| **Shared feeder wagon** | No | 48 (38.7%) |
|  | Yes | 76 (61.3%) |
| **Average (median) number of entries of the feeder wagon per week, when shared** | 7.1 (7) |  |
| **Shared machinery** | No | 60 (47.5%) |
|  | Yes | 64 (52.5%) |
| **Shared manure vehicle** | No | 103 (80.1%) |
|  | Yes | 21 (18.9%) |
| **Average (median) number of entries of the manure vehicle per month, when shared** | 0.39 (0.17) |  |
| **Shared materials (ear tag applicators, calving materials, cleaning materials or others)** | No | 100 (80.3%) |
|  | Yes | 24 (19.7%) |
| **Shared slurry vehicle** | No | 82 (66.2%) |
|  | Yes | 42 (33.8%) |
| **Average (median) number of entries of the slurry vehicle per month (when shared)** | 0.33 (0.08) |  |
| **Carcass deposit area** | Outside farm perimeter | 27 (20.5%) |
|  | Inside farm perimeter | 97 (79.5%) |
| **Average (median) number of entries of the carcass vehicle per month inside the farm perimeter (when applicable)** | 0.34 (0.17) |  |
| **Vehicle (slaughter/feedlot) may arrive carrying animals from outside the farm** | No | 5 (3.3%) |
|  | Not know | 5 (4.1%) |
|  | Yes | 114 (92.6%) |
| **Vehicles (slaughter/feedlot) may enter inside farm perimeter** | No | 6 (4.9%) |
|  | Yes | 118 (95.1%) |
| **Average (median) number of entries of the slaughter/feedlot vehicle per month inside farm perimeter** | 2.6 (2) |  |

Control of visits and staff also showed room for improvement (Table 4). Most farms had no perimeter fences (89.3%), and visitors’ parking was usually inside the farm perimeter (96.7%). Farms also commonly received visitors who, without protective clothing, had contact with the animals (92.6%).

**Table 4.** Biosecurity measures related to visitors and staff in 124 farms in Spain

|  |  | **N (%)** |
| --- | --- | --- |
| **(External) employees** | No | 68 (55.7%) |
|  | Yes, but do not work in other farms | 44 (34.4%) |
|  | Yes, and they work in other farms | 12 (9.8%) |
| **Perimeter fence** | Yes, always close | 5 (4.1%) |
|  | Yes, not always close | 8 (6.6%) |
|  | No | 111 (89.3%) |
| **Vehicle parking for visitors** | Outside farm perimeter | 4 (3.3%) |
|  | Inside farm perimeter | 120 (96.7%) |
| **Average (median) number of visitors per month that can contact animals** | 7.2 (5) |  |
| **Visitors use protective clothing** | Yes | 9 (7.4%) |
|  | No | 115 (92.6%) |

The MCA with standardized data explained 38.9% of the variance in biosecurity and health status among the 124farms. The percentage of variance explained by the first dimension was 24.7%, and for the second dimension was 14.2%.

The main results of the MCA are presented graphically in Fig 1. The object scores obtained from the MCA, together with the solutions of the TSCA, are presented in Fig 2. Three main clusters were formed that included 122 out of the 124 herds. S1 Table (within-cluster percentages) shows how each sanitary category or biosecurity variable is split within each cluster.

**Fig 1.**
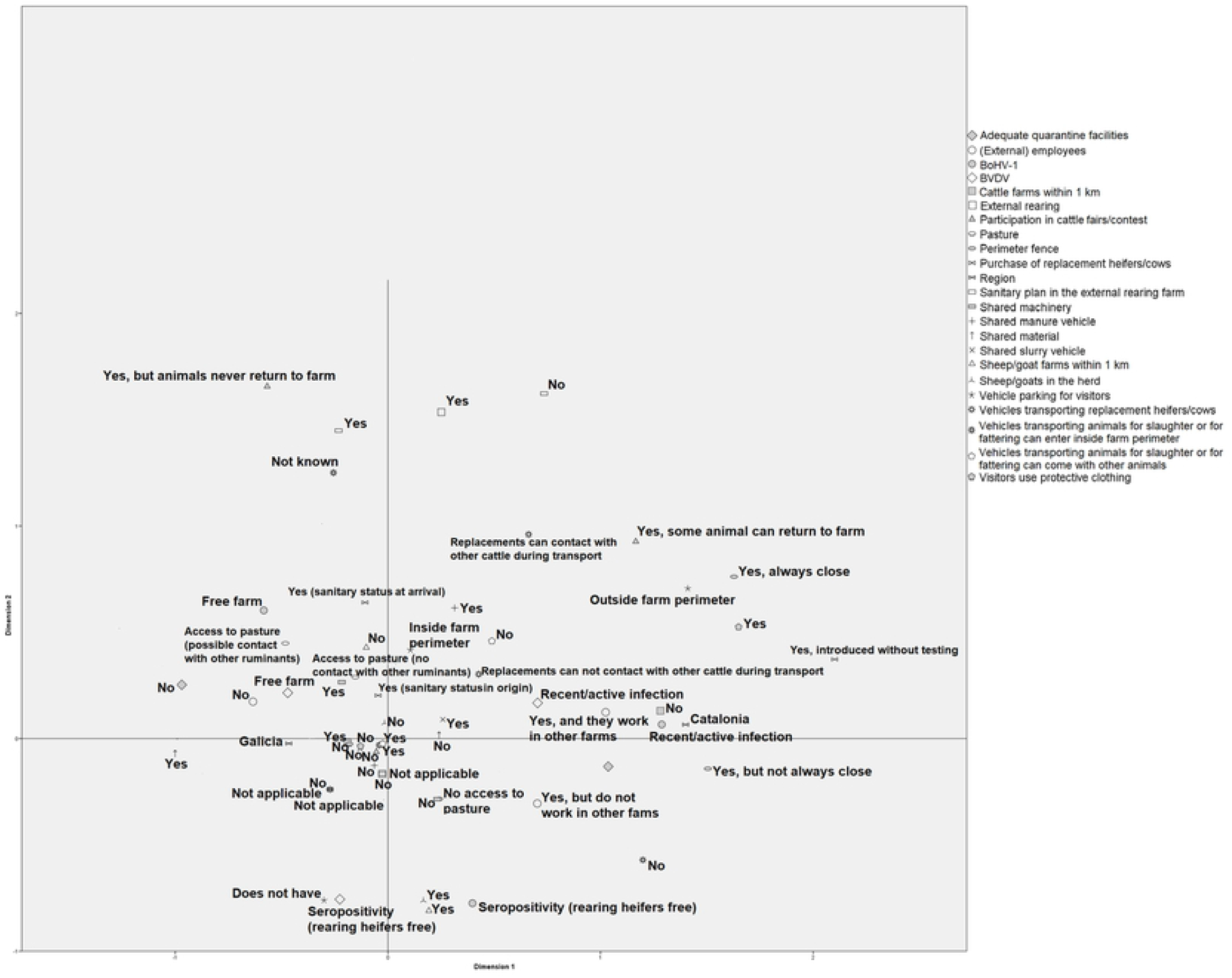
Joint MCA plot of category points for the different biosecurity measures, and BVDV/BoHV-1 profiles resulted from multiple correspondence analysis. Frequencies of cattle purchases, and entries of vehicles and visitors, each with 4 categories based on the quartiles of these frequencies, are not shown.

**Fig 2.**
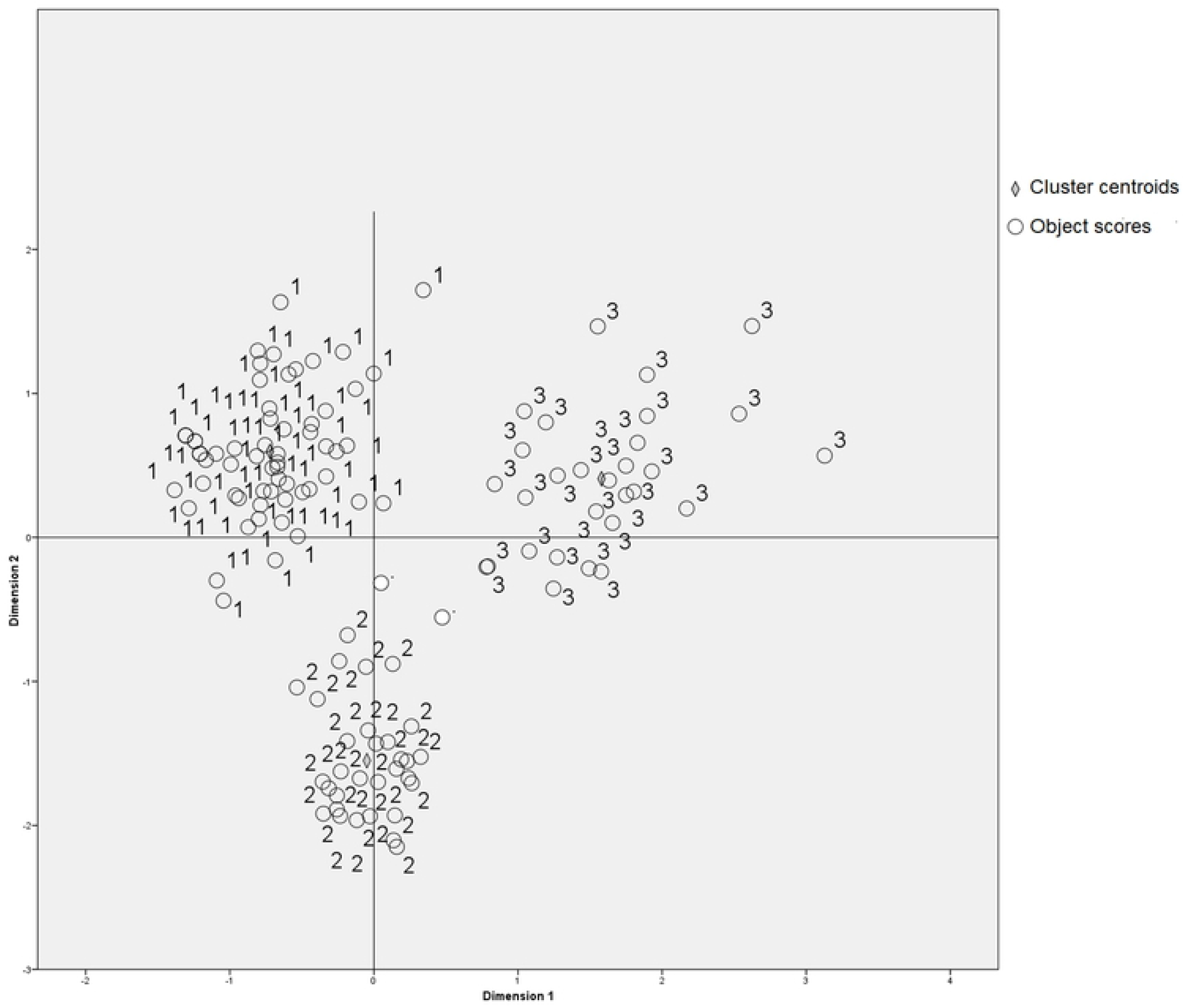
Object scores of the multiple correspondent analysis, and two-step cluster solution identifying 3 main clusters including 122 of the 124 study herds; the remaining 2 are shown in white in the center of the chart.

Cluster 1 (n=62) comprised herds located mainly in the upper left quadrant of the MCA chart. All these herds were from Galicia, with low mean herd size (51 cows). These farms were most frequently BVDV and BoHV-1-free.

These farms always checked the sanitary status of purchased animals, either at the farm of origin or on arrival. However, they did not have adequate quarantine facilities. This cluster was the one that most often used external rearing farms. Grazing was observed more frequently than in the other clusters, including possible contact between the farm’s cattle and ruminants from other farms. In addition, these herds were located in high-density cattle areas, with most located within 1 km of other cattle farms.

This cluster was also the one that most frequently shared machinery and materials with other farms.

The presence of external workers on Cluster 1 farms was very rare. Farms mostly lacked structures such as perimeter fencing or outdoor parking. Interestingly, these farms received numerous visitors despite their small size. Although the use of suitable protective clothing emerged as deficient in the entire sample of farms, it was especially inadequate in this cluster.

Cluster 2 (n=31) included herds also from Galicia, often of intermediate size (mean 63 cows). From the sanitary point of view, these farms often had seropositive animals but without evidence of recent or active infection.

These farms purchased animals less frequently than did those in other clusters. When they did so, purchases often entailed few animals that were tested, usually at their origin, against BVDV and BoHV-1. This was also the cluster with more control over the transport of purchased animals, avoiding contact with external cattle during transport. As in the previous cluster, these farms were located in high-density cattle areas, and most also had other small ruminant farms within 1 km.

Structures such as perimeter fencing or outdoor parking for visitors were also scarce in cluster 2 farms. Numbers of external workers were intermediate between those for clusters 1 and 3. Although cluster 2 farms received less visitors, the use of adequate protective clothing was still scarce.

Cluster 3 (n=29) comprised all herds from Catalonia, with the largest herds (mean 122 cows). This cluster had the highest proportions of active/recent infections with BVDV and BoHV-1.

Purchase of replacement animals was most common in cluster 3, and the number of purchased animals was usually high. Notably, the 11 farms that purchased animals without any testing were included in this cluster. Although infrequent, it was also the cluster with the highest frequency of participation in cattle fairs or contest.

Farms in this cluster shared machinery or materials less frequently than did the smaller farms in clusters 1 and 2,but manure vehicles were shared much more often than in the other two clusters. The frequency of entry of manure and slurry vehicles was also highest in cluster 3 farms, but shared use of feeder wagons was less frequently observed than in the other two clusters.

Structures such as perimeter fencing or outdoor parking for visitors were scarce, but nonetheless more frequent than in the other clusters. All cluster 3 farms employed external workers who often worked also on other farms. The use of protective clothing by visitors was somewhat more common in this cluster.

## Discussion

Spain is the eighth-largest dairy producer in the E.U., being responsible for 4.5% of the milk produced (19). Despite this, published information about biosecurity on Spanish dairy cattle farms is very scarce, and official or private initiatives for the implementation of biosecurity programs in this livestock sector are few.

Our study showed important short comings in the application of biosecurity measures in this sector, particularly highlighting room for improvement of such measures controlling various potential routes of disease introduction. Fortunately, most farms do not purchase animals, but among those who do there are still farms that do not test these animals. In addition, purchased cattle are habitually permitted to contact other cattle during transport process. Several types of vehicles, especially feeding vehicles, frequently enter farms. Policies related to visitors should also be improved: for example, nearly 93% of visitors that have contact with the animals do not use protective clothing.

A similarly lack of implementation of biosecurity measures has been observed on dairy farms elsewhere in Europe (7,11,20). The implementation of biosecurity plans on dairy farms is voluntary in almost all countries, with the exception of larger dairy farms in Denmark (21). Reg. (EU) No. 429/2016, which shall apply from 2021 and will affect EU animal health legislation, recognizes and addresses the importance of biosecurity (22). Therefore, farmers need to be motivated both to change existing behaviors, and to implement effective biosecurity practices to reduce the risk of disease introduction (23).

Our MCA and TSCA analyses show that there are different farm-typologies in relation to the implementation of biosecurity measures and their health status.

Cluster 1 includes farms with small herds and low turnovers that would pose a challenge to investment in infrastructure such as perimeter fencing or parking. For the same reason, such farms may frequently be forced to share machinery or materials. Another important point is that these small farms nonetheless receive a high number of visitors who come into contact with animals, both professional (veterinarians, technicians, etc.) and non-professional (neighboring farmers, etc.). Small farms more frequently require timely collaboration from neighboring farmers, for example, in the case of a calving, or receive courtesy visits from them, since such farms function with close links between the farm and the farmer’s own house, as has been described elsewhere (24). Despite the abovementioned lack of biosecurity measures, the proportion of such farms with recent/active BoHV-1 or BVDV infection was very low, possibly because of several factors.

One important factor is the nature of BVD and IBR control programs in Spain: these are voluntary, and executed only in some AC. In Galicia and other regions in the north-west and the Cantabric area, such programs are conducted mainly through Livestock Health Defense Associations (known as ADSG in Spanish) established by the regional governments in 2004.In Catalonia, these programs are absent. Farms included in cluster1 (and in cluster 2) were all from the Galicia region. Currently, ADSG groups 55.2% of the herds and 65.0% of Galician bovine census (12) and therefore are under control programs. For farms participating in ADSG, it is compulsory to test all purchased animals against BVDV and BoHV-1using established protocols (25). In the dairy cattle sector, the presence of the ADSG has increased awareness of the importance of incorporation of biosecurity controls into farming practices. Many farms that do not belong to ADSG now also practice such controls. Purchased cattle are considered to be the main risk factor for disease entry to dairy farms (3,26–29).

In contrast, cluster 3 included the largest herds in our study, located in Catalonia. Herd size has been previously described as a cluster variable for several biosecurity risks such as increased purchase of animals, increased visitors (veterinary practitioners, technicians), and the presence of external workers, all of which will increase the likelihood of disease introduction and maintenance (30). Cluster 3 farms in our study met all these characteristics, and, in addition, sometimes introduced animals without testing. Thus, although some facilities (i.e. perimeter fencing, outdoor parking) or biosecurity measures (use of protective clothing by visitors or scarce use of shared material or equipment) are more frequent in these farms, the cluster 3 showed the poorest sanitary level.

On the other hand, when comparing clusters 2 and 1, we observed that farms that purchase animals are somewhat more numerous in cluster 1, so herd size and number of animal purchases are not directly related in these two clusters. The smallest farms sometimes combine dairy production with other professional activities, using dairy farming to supplement family income (31). Thus, the lack of labor and even facilities for rearing heifers could explain the higher frequency of purchases than larger farms in cluster 2. Cluster 1 farms also use external rearing farms more frequently than do farms in cluster 2.

The most frequent sanitary status for cluster 2 was the presence of seropositive animals without evidence of recent or active infection.

The farms of cluster 2 are located in areas of high cattle density and very often close to small ruminant farms. In Spain, the particular case of BVDV circulation in sheep herds has been described (32,33);owners of these herds almost never implement BVDV and border disease virus control programs, and act only in cases of clinical outbreaks. Thereby, the risk they could pose is unknown.

The influence of our methodology on these results must be considered. Questionnaires were completed during face-to-face interviews on farms. This enabled us to explain the questions clearly, and to control for the bias related to social desirability response; in addition, the veterinarian responsible for the sanitary program of the farm was present. However, it is important to keep in mind that farmers enrolled voluntarily in the study, and therefore our results cannot be extrapolated directly to all dairy farms in Spain due to possible selection bias. Farmers more concerned with biosecurity might have decided to participate in the project, resulting in overrepresentation of farms with relatively good implementation of biosecurity measures in our sample. Additionally, the existence of voluntary ADSG BoHV-1 and BVDV control programs in Galicia but not in other regions may have reduced the representative value of our sample.

Nevertheless, despite the inherent limitations of this study we believe it to have provided a comprehensive overview of the main biosecurity shortcomings in the dairy sector of Spain. Such data should be useful to focus future training and to improve risk reduction strategies in this economically important industry.

## Acknowledgements

The authors want to thank all the farmers and veterinarians involved in the study. This research was supported by a project from the Ministry of Science and Innovation of Spain (AGL2016-77269-C2-2-R).

## Supporting information

**S1 Table. Composition of sanitary and biosecurity profiles within 3 clusters obtained through a two-step cluster analysis**

